# RAPTOR-targeted Antisense Oligonucleotides Restrict Pathological mTORC1 Activity in GATOR1-dependent Disease

**DOI:** 10.64898/2026.09.30.755052

**Authors:** Imane Hadj-Aissa, Maéline Muller, Antoine Desmeules, Laetitia Marcadet, Tristan Lachance-Brulotte, Nadine Morin, Marc-Étienne Huot, Jorge Soliz, Chantelle F. Sephton, Paul A. Dutchak

## Abstract

Loss-of-function mutations in the GATOR1 complex (NPRL2, NPRL3, and DEPDC5) are among the most common genetic causes of epilepsy. These mutations impair amino acid-dependent regulation of mTORC1 and cause hyperactive signaling, neuronal hyperexcitability, astrocytic metabolic remodeling, seizures, and early mortality. Here, we report the development of antisense oligonucleotides (ASOs) targeting RAPTOR, an mTORC1 scaffolding subunit, to correct pathological mTORC1 hyperactivity caused by defective upstream regulation. RAPTOR-targeted ASO treatment reduced RAPTOR expression in mouse and human systems and dampened mTORC1 signaling in GATOR1-deficient models. ASO-mediated RAPTOR reduction rescued neuronal hyperexcitability, corrected astrocytic mitochondrial dysfunction, and suppressed spontaneous seizures *in vivo*. A single treatment significantly extended longevity across neuronal and astrocytic models of GATOR1 deficiency. Collectively, these findings establish RAPTOR-targeted ASOs as a mechanism-based therapeutic approach that durably corrects cellular and circuit-level consequences of GATOR1 deficiency. By tuning RAPTOR abundance, this strategy provides a therapeutic framework for mTORC1-driven epilepsies and related diseases.

## Introduction

GATOR1 (GAP Activity Toward Rags 1) is an evolutionarily conserved amino acid-dependent signaling complex that inhibits the mechanistic target of rapamycin complex 1 (mTORC1) in response to amino acid deficiency (Bar-Peled et al., 2013; Loissell-Baltazar and Dokudovskaya, 2021; Neklesa and Davis, 2009). Loss-of-function mutations in GATOR1 subunits, including NPRL2 (Nitrogen Permease Regulator-like 2), NPRL3, and DEPDC5 (DEP domain-containing protein 5), are among the most common genetic causes of focal epilepsy and have also been reported in individuals with autism spectrum disorder (ASD) (Baldassari et al., 2019; Iffland et al., 2019; Ishida et al., 2013; Ricos et al., 2016). Epilepsy and ASD are frequently comorbid, with approximately 10% of individuals with ASD experiencing recurrent seizures, underscoring shared neurodevelopmental and circuit-level vulnerabilities (Liu et al., 2022). At the molecular level, genetic studies across both conditions increasingly converge on dysregulated mTORC1 signaling as an important pathogenic mechanism (Macnee et al., 2023; Moloney et al., 2021; Rolland et al., 2023). This convergence extends beyond GATOR1, as mutations affecting distinct upstream regulators of mTORC1, including TSC1/2 (Tuberous Sclerosis Complex) and PTEN (Phosphatase and Tensin homolog), similarly contribute to neuronal hyperexcitability and seizures, reinforcing mTORC1 as a convergent molecular node linking these conditions (Li et al., 2025; Winden et al., 2026). Importantly, sustained mTORC1 activation extends beyond its control of protein synthesis and cellular growth to reshape cellular metabolism and mitochondrial function, contributing to altered oxidative phosphorylation and increased oxidative stress in neurological mTORopathies (Hadj-Aissa et al., 2026; He et al., 2025; Li et al., 2026; Romagnolo et al., 2024).

Mechanistically, GATOR1 restrains amino acid-dependent mTORC1 activity by functioning as a GAP (GTPase-activating protein) for RAG GTPases, thereby limiting RAG-mediated recruitment of mTORC1 to the lysosomal surface and subsequent pathway activation (Bar-Peled et al., 2013). Loss of GATOR1 function disrupts this regulatory process, altering the normal integration of amino acid- and phosphatidylinositol 3-kinase (PI3K)-dependent signals that regulate mTORC1 activation (Bar-Peled et al., 2013; Muller et al., 2024; Shen et al., 2018). Central to this mechanism is RAPTOR (Regulatory-Associated Protein of mTORC1), a complex-defining scaffolding subunit that mediates mTORC1 recruitment to RAG GTPases at the lysosome in response to amino acid availability (Sancak et al., 2008). Genetic studies have shown that partial reduction of RAPTOR expression in heterozygous mice is sufficient to limit pathological mTORC1 signaling and improve neurological phenotypes in animal models of TSC (Karalis et al., 2022). These findings help establish RAPTOR dosage as an important determinant of pathway output and disease outcomes, raising the possibility that therapeutically tuning RAPTOR expression, rather than imposing broad and sustained kinase inhibition with small molecules, may provide a more precise and tolerable strategy for modulating mTORC1 activity *in vivo*.

Pharmacological inhibition of mTORC1 has demonstrated therapeutic potential in genetic epilepsies driven by mTORC1 hyperactivity. Studies have shown that small-molecule inhibitors of mTORC1 like rapamycin can reduce seizure burden and improve neurological outcomes in TSC (Franz et al., 2014; Franz et al., 2018; Meikle et al., 2008; Zeng et al., 2008), with similar benefit observed in GATOR1-deficient animal models (Ishida et al., 2022; Yuskaitis et al., 2019). These small-molecule inhibitors, however, have important limitations for long-term neurological use, including variable penetration into the central nervous system and reliance on repeated systemic dosing to sustain pathway suppression. Chronic systemic administration can produce adverse metabolic effects, including impaired glucose tolerance, increased hepatic gluconeogenesis, and insulin resistance (Houde et al., 2010). These observations indicate that sustained systemic inhibition of mTORC1 can disrupt whole-body metabolic homeostasis, raising concerns regarding the long-term tolerability and safety of chronic mTORC1 inhibition in early-onset genetic disease (Goul et al., 2023; Mannick and Lamming, 2023). These considerations highlight the need for therapeutic strategies that can normalize mTORC1 activity selectively within affected tissues while avoiding prolonged systemic pathway inhibition.

mTORC1 signaling is intrinsically dynamic, integrating nutrient availability, growth-factor cues, and cellular context to generate tightly regulated outputs over time (Sparta et al., 2023; Valvezan and Manning, 2019). Small-molecule inhibitors of mTORC1 can override these endogenous control mechanisms, causing pathway activity to become shaped by drug exposure and pharmacokinetics rather than physiological nutrient or growth-factor inputs. Consequently, small-molecule inhibition may perturb the graded and temporal responsiveness required for physiological mTORC1 regulation, particularly within the central nervous system, where mTORC1 influences neuronal excitability and metabolic coupling between cell types (Hadj-Aissa et al., 2026; He et al., 2025; Lipton and Sahin, 2014). We therefore hypothesized that durable correction of mTORC1-driven disease may benefit from therapeutic strategies that dampen pathological signaling capacity while preserving physiological responsiveness.

To address the limitations of small-molecule mTORC1 inhibitors, we pursued an RNA-based strategy to modulate mTORC1 activity through targeted reduction of RAPTOR expression. We conducted a systematic antisense oligonucleotide (ASO) screen targeting mouse and human *Rptor/RPTOR* mRNA to identify candidates capable of dampening mTORC1 signaling capacity rather than strictly inhibiting it. Lead ASOs were optimized through dose-response profiling and mechanistic validation in mouse and human cells prior to *in vivo* evaluation. Using NPRL2-deficient models of GATOR1-deficient epilepsy, RAPTOR-targeted ASO treatment corrected key phenotypic consequences of mTORC1 hyperactivity, including neuronal hyperexcitability and astrocytic reactivity. At the organismal level, ASO-mediated RAPTOR reduction suppressed spontaneous seizures and markedly extended longevity. Collectively, our results show that RAPTOR-targeted ASOs can durably dampen pathological mTORC1 hyperactivity and correct key disease-associated phenotypes without imposing blunt mTORC1 inhibition.

## Results

### Identification and validation of RAPTOR-targeting antisense oligonucleotides

To modulate the capacity of mTORC1 signaling at a structural level, we developed and screened antisense oligonucleotides (ASOs) targeting the mRNA of RAPTOR, a complex-defining scaffold protein of mTORC1. A panel of ASO molecules was designed to target sites across the *Rptor/RPTOR* mRNA in both mouse and human transcripts excluding any sequences with homology to off-target transcripts as determined using NCBI BLAST analysis. Candidate ASOs were evaluated by their ability to repress RAPTOR expression in mouse embryonic fibroblasts (MEFs) and astrocytic cells (C8-D1A). Screening was initially performed using MEF cells treated with 5 µM of each candidate ASO or infected with a lentivirus expressing RAPTOR shRNA as a positive control (Figure 1A). RAPTOR reduction was observed using western blot analysis following treatment with ASO2 or ASO3 in both MEF and C8-D1A cells (Figure 1A, 1B). Based on these results and its proximal location relative to the translational start site, ASO2 was selected for further dose-response analysis against a control scrambled-ASO2 (scrASO2), generated by rearranging the position of four nitrogenous bases within the ASO2 sequence. NCBI BLAST analysis also confirmed that scrASO2 showed no full-length match to any annotated transcripts. ASO2 treatment of MEF cells produced a dose-dependent reduction of RAPTOR compared to scrASO2, reaching statistical significance from 0.5 to 5 µM following 48 hours of treatment (Figure 1C). A tiling approach was also used to evaluate the efficacy of RAPTOR knockdown by targeting conserved sequence regions around ASO2 in both MEF and human HEK293 cells; however, the initial ASO2 sequence produced the strongest reduction of RAPTOR expression in these assays (Figure 1D). Next, we used quantitative reverse transcription-PCR (QRT-PCR) to determine if ASO-mediated reduction of *Rptor/RPTOR* mRNA occurred in MEF or HEK293 cells, comparing scrASO2 and ASO2 after 72 hours of treatment. Consistent with protein level decreases, QRT-PCR analysis also showed a significant reduction of *Rptor/RPTOR* mRNA in both species (Figure 1E). To verify that ASO2-mediated RAPTOR reduction was independent of specific genetic defects in upstream regulators of mTORC1, we treated several MEF cell lines expressing epilepsy-linked mutations in NPRL2, as well as NPRL3 KO cells, previously generated (Muller et al., 2024). Western blot analysis confirmed that ASO2 treatment reduced RAPTOR expression in these cells, independent of the upstream genetic defect (Figure 1F). Collectively, these data show that ASO2 treatment efficiently reduces the mRNA and protein expression of RAPTOR in both mouse and human cells.

**Figure 1.**
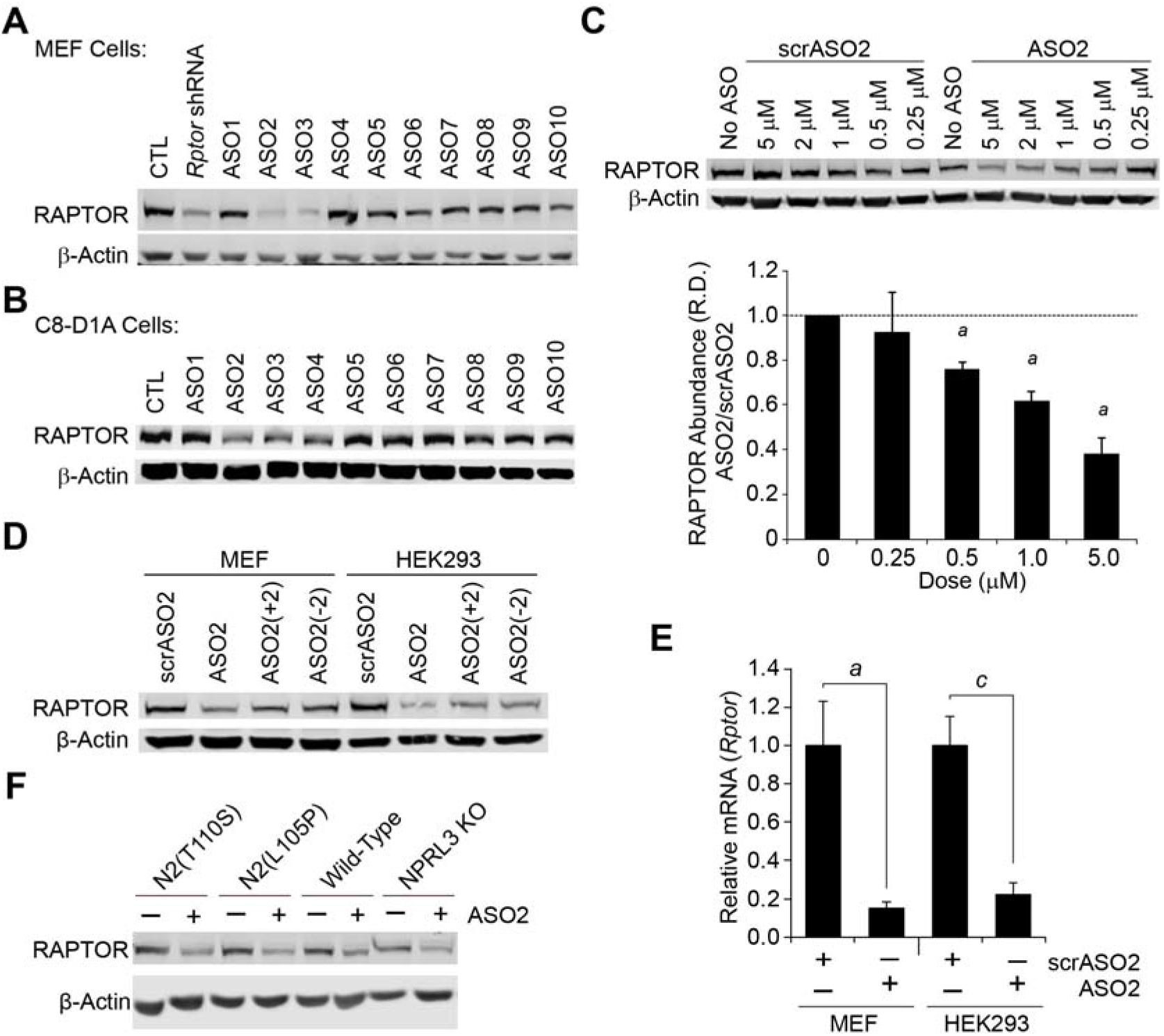
Identification and validation of RAPTOR-targeted antisense oligonucleotides. (A)Screen of RAPTOR-targeted antisense oligonucleotides (ASO1-ASO10) in MEF cells, with RAPTOR shRNA-expressing lentivirus included as a positive control. Cells were treated with 5 µM ASO or transduced with lentivirus for 48 hours, followed by western blotting against RAPTOR and β-actin as an internal control. (B)Validation screen of ASO1-ASO10 in C8-D1A astrocytes. Cells were treated with 5 µM ASOs for 72 hours followed by western blotting against RAPTOR and β-actin as an internal control. (C)Dose-response analysis of ASO2 compared to scrambled control (scrASO2) in MEF cells treated for 48 hours. Protein levels were assessed by western blotting against RAPTOR and β-actin as an internal control. RAPTOR signal intensity was normalized to β-actin and expressed as relative density (R.D.) compared to scrASO2 from three independent experiments. (D)Tiling analysis of ASO2 sequence variants (±2 nucleotide shifts) in MEF and HEK293 cells treated with 5 µM of each ASO for 48 hours. Representative western blots of RAPTOR and β-actin from three independent experiments are shown. (E)Quantitative RT-PCR analysis of *Rptor/RPTOR* mRNA expression in MEF and HEK293 cells treated with 5 µM ASO2 for 72 hours; U36B4 used as an internal control. Data represent three independent cultures per condition, analyzed in triplicate reactions. (F)Western blot analysis of RAPTOR protein expression in GATOR1-deficient MEF models (NPRL2-T110S, NPRL2-L105P, and NPRL3 KO) following 5 µM ASO2 treatment for 48 hours. Quantitative data are shown as the mean ± SEM. Statistical significance was determined using two-tailed Welch’s *t*-test; *a*, p<0.05 versus scrASO2; *c*, p<0.001 versus scrASO2.

### RAPTOR-targeted ASO2 achieves *in vivo* target engagement and therapeutic benefit across neuronal and astrocytic models of GATOR1 deficiency

To determine whether RAPTOR-targeted ASOs could reduce RAPTOR *in vivo*, we examined RAPTOR expression in the adult mouse brain 14 days after bilateral stereotactic intracerebroventricular (ICV) administration of ASO2 (100 µg per hemisphere) or sham treatment. ASO2 significantly reduced RAPTOR expression across the examined brain regions, as determined by immunohistochemistry (IHC) and confocal microscopy with tiled image acquisition (Figure 2A). Quantification of the mean RAPTOR fluorescence intensity within the tissue showed a ∼50% reduction relative to control (Figure 2B). RAPTOR reduction was also independently confirmed using western blot analysis in a separate cohort following ICV injection of ASO2 or sham control (Figure S1A, S1B). To determine whether ASO2 could functionally dampen GATOR1-dependent mTORC1 hyperactivation *in vivo*, we next treated NPRL2 astroKO mice (GFAP-IRES*-Cre; Nprl2*^*fl/fl*^) and littermate controls. Notably, NPRL2 astroKO mice show increased phosphorylation of downstream mTORC1 signaling, including ribosomal protein S6 (Hadj-Aissa et al., 2026). Mice were treated at 4 months of age using bilateral stereotactic hippocampal injections of ASO2 (100 µg per hemisphere) or sham control, followed by tissue collection and western blotting one month later. Consistent with our IHC observations, ASO2 treatment significantly decreased RAPTOR expression and reduced P-S6 in control and NPRL2 astroKO brains to comparable levels, despite the elevated baseline in NPRL2 astroKO mice (Figure S1C-E). Collectively, these data show that ASO2 treatment durably represses RAPTOR expression and dampens hyperactive mTORC1 signaling in the brain.

**Figure 2.**
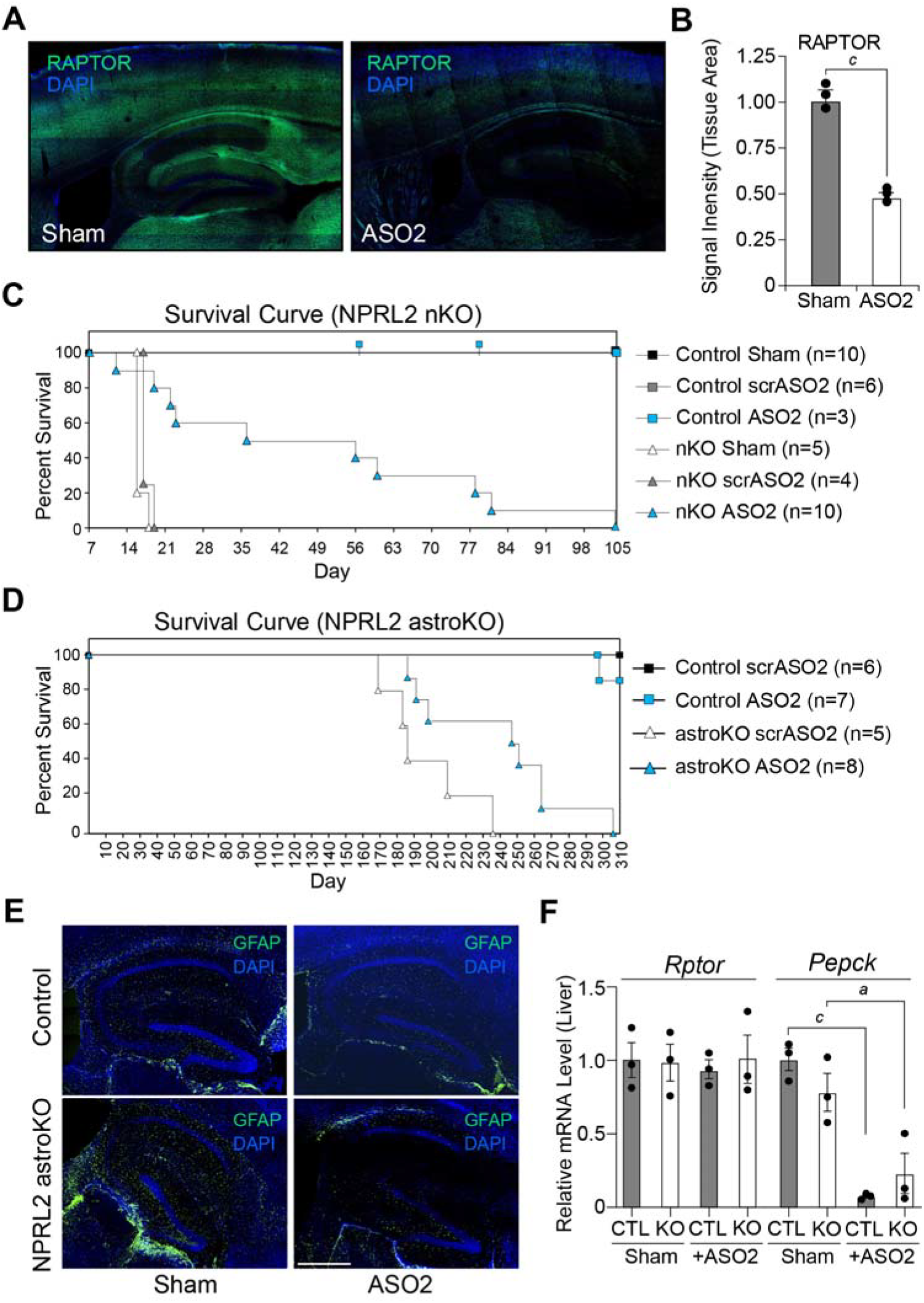
RAPTOR-targeted ASO2 reduces brain RAPTOR expression and improves disease outcomes in GATOR1-deficient mouse models. (A)3-month old adult mice received bilateral stereotactic ICV injections with 100 µg/side ASO2 (PBS vehicle) or sham control and were harvested 14 days later. Brain sections were analyzed by confocal microscopy with tiled image acquisition for RAPTOR (green) and DAPI (blue). Representative images of three independent animals per condition. (B)Quantification of RAPTOR fluorescence signal intensity within the brain, normalized to tissue area defined by combined RAPTOR and DAPI signal. Data represent three independent animals per group. (C)Kaplan-Meier survival analysis of NPRL2 nKO and littermate controls following free-hand ICV treatment with 50 µg/side ASO2, scrASO2, or sham injection at postnatal day 5. Tick marks indicate censored control animals treated with ASO2. (D)Kaplan-Meier survival analysis of NPRL2 astroKO mice and littermate controls following hippocampal treatment with 100 µg/side ASO2 or scrASO2 injection at 5 months of age. (E)Immunohistochemistry and confocal microscopy images with tiled image acquisition of hippocampal sections from control and NPRL2 astroKO mice treated with sham or ASO2 at 4 months of age and harvested one month later, showing GFAP (green) and DAPI (blue). (F)Quantitative RT-PCR analysis of hepatic *Pepck* (phosphoenolepyruvate carboxykinase) and *Rptor* expression in control and NPRL2 astroKO mice following hippocampal ASO2 or sham treatment at 5 months of age and measured one month post-injection. Data represent three animals per group. Quantitative data are shown as the mean ± SEM. Statistical significance was determined using two-tailed Welch’s *t*-test; *a*, p<0.05 versus sham; *c*, p<0.001 versus sham.

Having observed a robust and sustained reduction of RAPTOR expression following ASO2 treatment *in vivo*, we asked whether ASO2 could ameliorate GATOR1-dependent disease outcomes. We evaluated RAPTOR-targeted ASO treatment using two distinct models of GATOR1-deficiency lacking NPRL2 expression either in excitatory glutamatergic neurons (Hui et al., 2022) or astrocytes (Hadj-Aissa et al., 2026). We first evaluated ASO2 in NPRL2 nKO mice, a severe model characterized by early-onset seizures and mortality before P21 (postnatal day 21). We treated mice once at P5 using free-hand ICV injections with either ASO2 or scrASO2 (50 µg per hemisphere), with sham-treated animals included as an additional control. In early cohorts, all treated animals were maintained until all NPRL2 nKO mice reached endpoint, after which matched ASO2-treated control mice were euthanized and were recorded as censored observations in the survival curve (tick marks, Figure 2C). All control-treated NPRL2 nKO mice died before P19, whereas ASO2 treatment markedly extended longevity, with 50% of treated NPRL2 nKO mice living beyond 55 days and one animal reaching P104 (Figure 2C). This represents a nearly three-fold increase in mean lifespan in this model (log-rank Mantel-Cox test, p<0.0001). Moreover, ASO2-treated NPRL2 nKO mice remained independently viable after weaning, demonstrating a substantial rescue beyond longevity alone.

To determine whether improved viability was accompanied by a suppression of seizures, additional cohorts of ASO-treated control and NPRL2 nKO mice were video-recorded for 5 weeks after weaning. Video analysis revealed a marked suppression of seizure activity following ASO2 treatment, with 22% of the ASO2-treated NPRL2 nKO mice experiencing seizure episodes one week after weaning and no additional seizures detected during the following 2 to 5 weeks of monitoring (Table 1). We next assessed whether the sustained reduction of RAPTOR in the brain produced any peripheral hematological abnormalities by performing complete blood counts (CBC) in animals harvested at P51-54. ASO2-rescued NPRL2 nKO mice showed no evidence of leukopenia, granulocytopenia, lymphopenia, eosinophilia, or anemia relative to scrASO2 or ASO2-treated control mice (Table 2). Given that thrombocytopenia is a recognized hematological concern for some ASO-based approaches, we also examined platelet counts in the panel. Platelet counts were modestly lower in ASO2-treated NPRL2 nKO mice relative to scrASO2-treated controls, without a corresponding significant change in mean platelet volume (MPV). Modest changes in red-cell indices were also observed, including a reduced red blood cell distribution width (RDW) and increased mean corpuscular hemoglobin concentration (MCHC) in ASO2-treated NPRL2 nKO mice compared to scrASO2-treated controls. Together, these data show that ASO-mediated RAPTOR reduction achieves sustained suppression of seizure activity and substantially extends longevity in severe GATOR1-dependent epilepsy, without evidence of overt hematological toxicity.

**Table 1.** Seizure episodes during weekly 24-hour video monitoring in control and NPRL2 nKO mice following P5 ICV ASO treatment with 50 µg/hemisphere. Mice were monitored for 5 consecutive weeks after weaning at P21. Values indicate the total number of seizure episodes detected during each weekly recording period. In the NPRL2 nKO + ASO2 group, 33 seizure episodes were detected in 2 of 9 mice during week 1. No seizure episodes were detected in either control group during the monitoring period.

| Seizure Episode Count |  |  |  |
| --- | --- | --- | --- |
| Week | Control + scrASO2<br>(n=10) | Control + ASO2<br>(n=10) | NPRL2 nKO +ASO2<br>(n=9) |
| 1 | 0 | 0 | 33 total (2/9 mice) |
| 2 | 0 | 0 | 0 |
| 3 | 0 | 0 | 0 |
| 4 | 0 | 0 | 0 |
| 5 | 0 | 0 | 0 |

**Table 2.** Complete blood count analysis of control and NPRL2 nKO mice at P51-54 following free-hand ICV treatment with 50 µg/hemisphere of scrASO2 or ASO2 at P5. Statistical comparisons were performed using two-tailed Welch’s unpaired *t*-tests. Data are shown as mean ± SD.

| Parameter | Control +<br>scrASO2<br>(Group A, n=10) | Control +<br>ASO2<br>(Group B, n=7) | N2 nKO +<br>ASO2<br>(Group C, n=7) | A vs B<br>p-value | A vs C<br>p-value | B vs C<br>p-value |
| --- | --- | --- | --- | --- | --- | --- |
| EOS# 10<sup>3</sup>/mm<sup>3</sup> | 0.13 ± 0.17 | 0.37 ± 0.36 | 0.21 ± 0.30 | 0.137 | 0.544 | 0.379 |
| GRA# 10<sup>3</sup>/mm<sup>3</sup> | 1.02 ± 0.43 | 1.43 ± 0.75 | 1.00 ± 0.89 | 0.224 | 0.957 | 0.348 |
| LYM# 10<sup>3</sup>/mm<sup>3</sup> | 6.70 ± 1.66 | 6.67 ± 2.66 | 6.50 ± 3.56 | 0.980 | 0.894 | 0.921 |
| MON#<br>10<sup>3</sup>/mm<sup>3</sup> | 0.18 ± 0.08 | 0.19 ± 0.09 | 0.14 ± 0.14 | 0.894 | 0.541 | 0.510 |
| PLT# 10<sup>3</sup>/mm<sup>3</sup> | 903.60 ±<br>113.02 | 752.29 ±<br>221.39 | 632.43 ±<br>231.47 | 0.134 | **0.021** | 0.342 |
| MPV µm<sup>3</sup> | 5.75 ± 0.31 | 5.74 ± 0.23 | 5.56 ± 0.28 | 0.957 | 0.199 | 0.197 |
| WBC# 10<sup>3</sup>/mm<sup>3</sup> | 7.90 ± 2.09 | 8.29 ± 3.43 | 7.64 ± 4.54 | 0.797 | 0.892 | 0.771 |
| RBC 10<sup>6</sup>/mm<sup>3</sup> | 11.44 ± 1.58 | 10.31 ± 1.32 | 9.56 ± 3.22 | 0.132 | 0.189 | 0.581 |
| RDW % | 15.80 ± 0.24 | 15.74 ± 0.34 | 15.49 ± 0.30 | 0.709 | **0.043** | 0.158 |
| HCT % | 53.01 ± 7.43 | 47.11 ± 6.05 | 43.41 ± 14.86 | 0.093 | 0.153 | 0.559 |
| HGB g/dL | 17.99 ± 2.46 | 15.99 ± 1.84 | 15.26 ± 4.89 | 0.074 | 0.209 | 0.722 |
| MCHC g/dL | 33.93 ± 0.60 | 33.99 ± 0.97 | 35.39 ± 1.47 | 0.895 | **0.040** | 0.061 |

Since ASO2 reduced RAPTOR expression and hyperactive mTORC1 signaling in the brain of NPRL2 astroKO mice (Figure S1C), we asked whether this molecular correction was associated with therapeutic benefit in this model, which develops reactive astrocytes, seizures, and early death (Hadj-Aissa et al., 2026). Here, NPRL2 astroKO and control mice received bilateral stereotactic hippocampal injections of ASO2 or scrASO2 (100 μg per hemisphere) at 5 months of age. ASO2 treatment significantly extended the longevity of NPRL2 astroKO mice, delaying the first death by 19 days and increasing the median lifespan from 185 days in scrASO2-treated animals to approximately 250 days, with one animal surviving to P306 (Figure 2D). Kaplan-Meier analysis confirmed a statistically significant longevity benefit following ASO2 treatment in the NPRL2 astroKO mice (log-rank Mantel-Cox test, p<0.01), consistent with the benefit observed in NPRL2 nKO mice (Figure 2C). Given the variable onset of spontaneous seizures and death of NPRL2 astroKO mice, we performed an exploratory video-monitoring study in a small cohort before and after ASO2 treatment. Seizures were observed in 2 of 4 animals during pretreatment monitoring, whereas no seizures were detected during a 48-hour video recording period one week after ASO2 administration (data not shown).

To investigate whether the longevity benefit of RAPTOR-targeted ASO treatment was accompanied by correction of astrocytic pathology, NPRL2 astroKO and control mice were treated at 4 months of age and analyzed one month later using immunohistochemistry and confocal microscopy. NPRL2 astroKO mice showed pronounced astrocytic reactivity, whereas ASO2 treatment markedly reduced GFAP immunoreactivity and attenuated reactive morphology across the examined tissue (Figure 2E, S2A-E). We next tested whether ASO2 could reduce the oxidative tissue damage associated with GATOR1 deficiency. 4-hydroxynonenal (4-HNE), a marker of lipid peroxidation, was significantly elevated in hippocampal sections from NPRL2 astroKO mice relative to controls, whereas ASO2 treatment significantly reduced 4-HNE signal intensity in these mice to control levels (Figure S2F, S2G). These data demonstrate that ASO-mediated RAPTOR reduction ameliorates both astrocytic reactivity and oxidative damage in the NPRL2 astroKO brain *in vivo*.

To test whether ASO2 treatment affected recognition memory, control and NPRL2 astroKO mice were treated with ASO2, scrASO2, or sham control, and subjected to a novel object recognition test one month after dosing. ASO2 treatment did not impair novel object recognition performance in either control or NPRL2 astroKO mice under the examined conditions (Figure S3A). To assess the potential for systemic metabolic effects following ASO treatment, particularly hyperglycemia associated with systemic mTORC1 inhibition (Houde et al., 2010), we also analyzed hepatic *Rptor* and *Pepck* mRNA expression in the mice one month after treatment. No difference in hepatic *Rptor* mRNA expression was observed between control and NPRL2 astroKO mice treated by hippocampal injection with either sham or ASO2 (Figure 2F). In contrast, hepatic *Pepck* expression was reduced to a similar extent in ASO2-treated control and NPRL2 astroKO mice, with no significant increase in blood glucose at either one week or one month post-treatment (Figure 2F, S3B). These results show that brain-directed RAPTOR-targeted ASO treatment does not impair recognition memory or induce hyperglycemia under the conditions examined. Combined with the effects on longevity, astrocytic reactivity, and oxidative damage, these data support a durable correction of mTORC1-driven pathologies across distinct animal models of GATOR1-dependent epilepsy, without evidence of impaired recognition memory or hyperglycemia under the conditions examined.

### RAPTOR-targeted ASO2 normalizes neuronal network hyperactivity

Having demonstrated therapeutic efficacy for ASO2 across neuronal and astrocytic models *in vivo*, we next asked whether RAPTOR reduction could directly ameliorate the neuronal hyperexcitability associated with GATOR1-deficiency. To investigate the impact of ASO-mediated RAPTOR reduction on neural network activity, we generated primary cortical neuron cultures using *Nprl2*^*fl/fl*^ mice as a model of GATOR1-dependent mTORC1 hyperactivity, as previously described (Hui et al., 2022). *Nprl2*^*fl/fl*^ cells were plated onto 24-well multi-electrode plates and infected with lentivirus expressing either Cre recombinase or a negative control to generate NPRL2 KO and control cultures. Cultures received a single treatment with either 0.5 µM ASO2 or scrASO2 at DIV4 (days *in vitro*; Figure 3A), and functional network activity was monitored using the Axion Biosystems Maestro multi-electrode array (MEA) system from DIV6 to DIV21.

**Figure 3.**
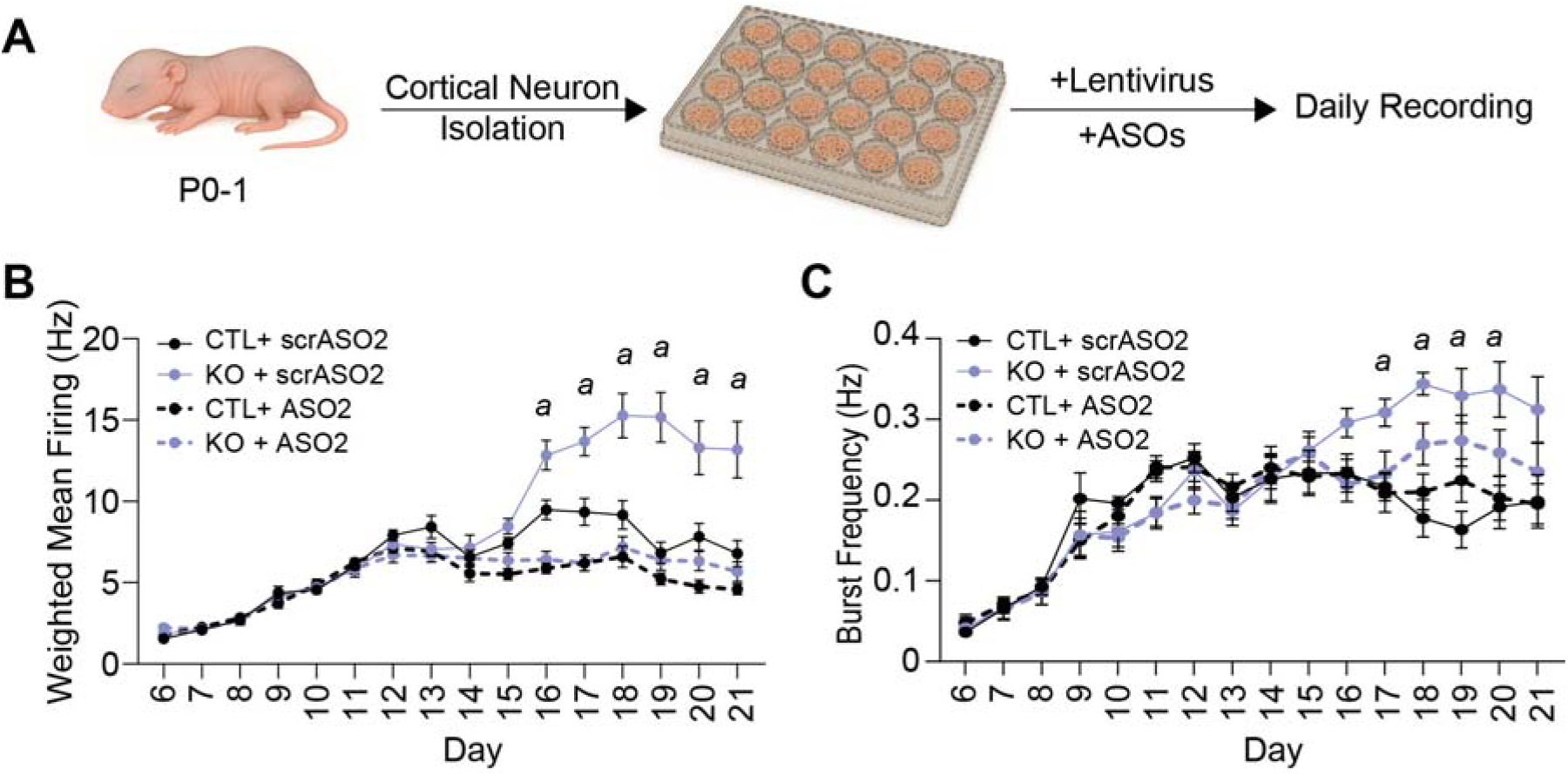
RAPTOR-targeted ASO2 normalizes network hyperactivity in NPRL2-deficient primary neuronal cultures. (A)Experimental schematic of primary cortical neuron cultures established from postnatal day 0-1 (P0-1) NPRL2^fl/fl^ pups and cultured on multielectrode array (MEA) plates. Neuronal cultures were transduced with lentivirus expressing Cre recombinase or control virus at DIV1, followed by treatment with 0.5 µM ASO2 or scrASO2 at DIV4. Cartoon images were generated using Microsoft Copilot. (B)Weighted mean firing rate (Hz) recorded using the Axion Maestro MEA system from DIV6 to DIV21. Data represent the mean ± SEM of five independent experiments with three wells per condition in each experiment. (C)Network burst frequency measured from DIV6 to DIV21 using the Axion Maestro MEA system. Data represent the mean ± SEM of five independent experiments with three wells per condition in each experiment. Quantitative data are shown as the mean ± SEM. Statistical significance was determined using two-tailed unpaired Welch’s *t*-test; *a*, p<0.05; scrASO2-treated KO versus scrASO2-treated control.

NPRL2 KO neuronal cultures developed robust network hyperexcitability, with approximately three-fold higher weighted mean firing rates from DIV16 through DIV21 compared to control cultures (Figure 3B). A single treatment at DIV4 with RAPTOR-targeted ASO2 durably normalized this hyperactivity and reduced firing rates to control levels (Figure 3B). NPRL2 KO cultures also showed an increase in burst frequency relative to controls, with significant increases from DIV17 to DIV20. ASO2 treatment similarly reduced burst frequency without suppressing activity below control levels (Figure 3C). These findings provide an *in vitro* functional correlate of the seizure suppression observed *in vivo*, and demonstrate that partial RAPTOR reduction can durably normalize pathological neuronal activity without producing a generalized network suppression.

### RAPTOR-targeted ASO2 corrects pathological astrocytic phenotypes across mouse and human systems

We previously reported that GATOR1-deficient astrocytes show elevated mitochondrial reactive oxygen species (ROS), electron transport chain dysfunction, and redox imbalance (Hadj-Aissa et al., 2026). To determine whether RAPTOR-targeted ASO treatment could correct these defects, we treated control and NPRL3 KO C8-D1A astrocytic cell lines with 1 µM of ASO2 or scrASO2 for 3 days, followed by MitoSOX Red staining and epifluorescence microscopy to visualize mitochondrial ROS production. MitoSOX fluorescence intensity was significantly elevated in NPRL3 KO cells compared with control, as previously reported (Hadj-Aissa et al., 2026). ASO2 treatment significantly reduced MitoSOX fluorescence in the NPRL3 KO astrocytes to levels in control cells (Figure 4A, 4B). To determine if this correction was accompanied by improvements in mitochondrial respiratory function, we next performed high-resolution respirometry using the Oroboros O2k instrument to assess electron transport chain activity. scrASO2-treated NPRL3 KO astrocytes showed a marked reduction in Complex II- and Complex IV-linked respiration compared to control cells, whereas ASO2 treatment of NPRL3 KO astrocytes corrected these respiration defects to control levels (Figure 4C). Together, these findings demonstrate that ASO-mediated RAPTOR reduction ameliorates mitochondrial and redox dysfunction in GATOR1-deficient astrocytes.

**Figure 4.**
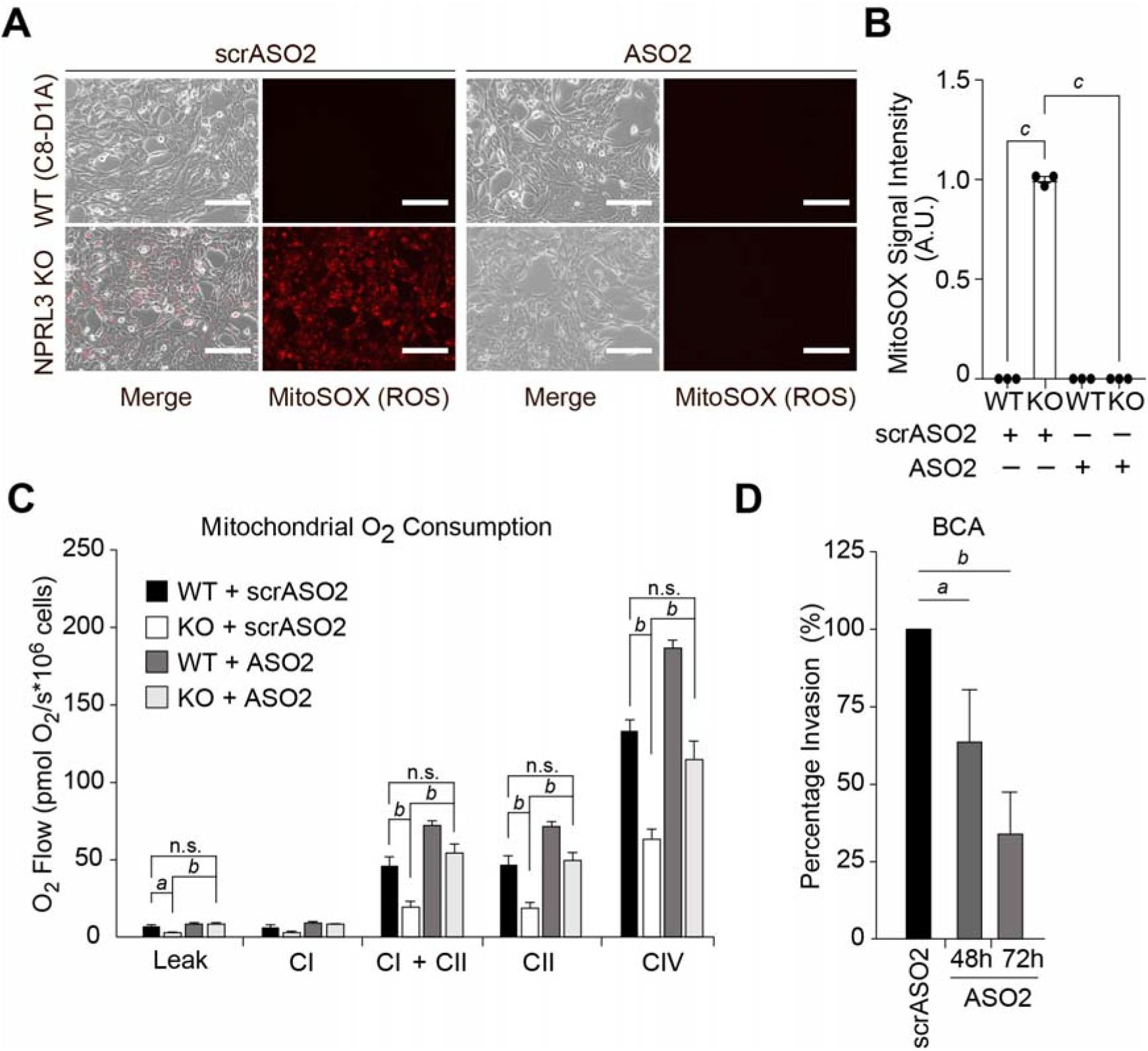
RAPTOR-targeted ASO2 reduces oxidative stress, restores mitochondrial respiration, and decreases human astrocyte invasion. (A)Representative epifluorescence microscopy images of MitoSOX Red staining in wild-type (WT) and NPRL3 knockout (NPRL3 KO) C8-D1A astrocytes treated with 1 µM ASO2 or scrASO2 for 72 hours. Data represent three independent experiments. (B)Quantification of mitochondrial reactive oxygen species (ROS) levels from (A) expressed as relative fluorescence intensity. Data represent three independent experiments. *c*, p<0.001. (C)Mitochondrial oxygen consumption rates (OCR) measured in WT and NPRL3 KO astrocytes following treatment with 1 µM ASO2 or scrASO2 for 72 hours, assessed by high-resolution respirometry. Data represent 3-independent experiments. (D)Boyden chamber invasion assay (BCA) in normal human astrocytes (NHA) treated with 1 µM ASO2 or scrASO2 for 48 or 72 hours. Invasion is expressed as relative percentage normalized to scrASO2-treated controls at each time point. Data represent 3-independent experiments. Quantitative data are shown as the mean ± SEM. Statistical significance was determined using two-tailed Welch’s *t*-test; *a*, p<0.05; *b*, p<0.01; *c*, p<0.001; n.s., not significant.

mTORC1 hyperactivity in astrocytes is associated with reactive morphological remodeling and spatial reorganization within epileptic tissue, phenotypes that can be accompanied by increased astrocytic invasiveness *in vitro* (Li et al., 2015). To determine whether RAPTOR-targeted ASO2 could modulate this phenotype in human cells, we assessed the invasiveness of normal human astrocytes (NHA cells) using a Boyden chamber assay. Cells were treated with 0.5 µM of either scrASO2 or ASO2 for 48 or 72 hours, followed by image-based quantification of invasiveness into the chamber. RAPTOR-targeted ASO2 treatment reduced astrocyte invasion at both time points relative to scrASO2-treated controls (Figure 4D). These findings demonstrate that ASO2 is functionally active in human astrocytes and can reduce astrocyte invasion. Collectively, these results show that RAPTOR-targeted ASO2 corrects mitochondrial and redox dysfunction in GATOR1-deficient astrocytes while also reducing an invasive phenotype in human astrocytes.

### RAPTOR-targeted ASO2 partially restores GATOR1-dependent nutrient-responsive mTORC1 signaling

Having established a functional rescue of GATOR1 deficiency in both neuronal and astrocytic systems following ASO2 treatment, we next asked whether partial RAPTOR reduction could restore the core cell signaling defect caused by GATOR1-deficiency. To address this, we used wild-type and NPRL2 KO MEF cells, a well-established model for assessing GATOR1-dependent amino acid sensing (Dutchak et al., 2015; Muller et al., 2024). Cells were treated with either scrASO2 or RAPTOR-targeted ASO2 for 3 days in complete media, followed by nutrient deprivation in Earle’s balanced saline solution (EBSS) for 1 hour. In control cells, EBSS treatment significantly reduced the phosphorylation of downstream mTORC1 targets, P-S6K (S6 Kinase) and P-S6, consistent with nutrient-responsive suppression of mTORC1 signaling (Figure 5A). In contrast, NPRL2 KO cells retained elevated levels of P-S6K and P-S6 during nutrient deprivation, consistent with an impaired GATOR1-dependent suppression of mTORC1. ASO2 treatment reduced RAPTOR protein expression in NPRL2 KO MEFs and shifted mTORC1 signaling toward a more nutrient-responsive state, with lower P-S6K and P-S6 levels following EBSS treatment, compared to scrASO2-treated NPRL2 KO cells (Figure 5B). Consistent with a partial relief of mTORC1/S6K-dependent negative feedback, ASO2-treated control and NPRL2 KO MEF cells also showed a modest increase in AKT phosphorylation at Ser473 in the fed and starved conditions, respectively. These finding demonstrate that partial reduction of RAPTOR can improve nutrient-responsive suppression of mTORC1 signaling despite the loss of GATOR1 function (Figure 5C).

**Figure 5.**
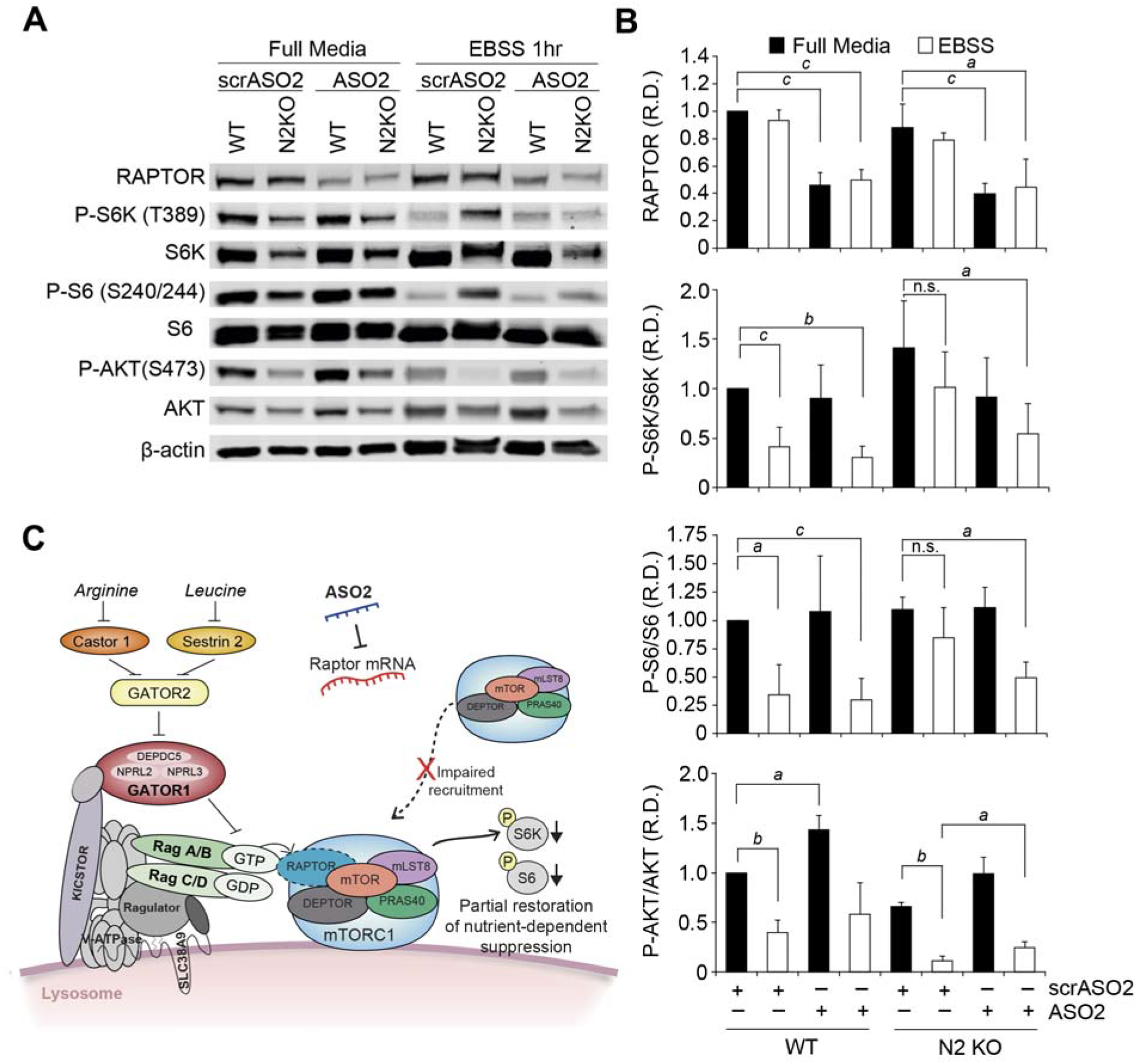
RAPTOR-targeted ASO2 partially restores nutrient-dependent regulation of hyperactive mTORC1-dependent cell signaling. (A)WT and NPRL2 KO MEF cells were treated with 5 µM ASO2 or scrASO2 for 72 hours and then either maintained in complete media or starved in EBSS for 1 hour prior to harvest. Protein lysates were analyzed by western blotting for P-S6K (T389), S6K, P-S6 (S240/244), S6, P-AKT (S473), AKT, RAPTOR, and β-actin. Date represent at least three independent experiments. (B)Quantification of western blot signal intensity from (A). Phosphorylated S6K, S6, and AKT signals were normalized to their respective total protein levels and RAPTOR abundance was normalized to β-actin. Date are expressed as relative density (R.D.) and represent three independent experiments. (C)Schematic model illustrating the proposed mechanism of RAPTOR-targeted ASO acting in mTORC1 signaling. Reduction of RAPTOR abundance limits mTORC1 recruitment to RAG GTPases at the lysosomal surface, thereby restricting pathway activation. This decreases phosphorylation of downstream targets and partially restores the suppression of mTORC1 signaling during nutrient deprivation. Quantitative data are shown as the mean ± SEM. Statistical significance was determined using two-tailed Welch’s t-test; a, p<0.05; b, p<0.01; n.s.=not significant.

## Discussion

In this study, we developed and characterized a novel antisense oligonucleotide strategy to reduce RAPTOR expression and dampen mTORC1 signaling capacity in mouse and human systems. Through systematic screening and validation, we identified several RAPTOR-targeted ASOs with robust activity and selected ASO2 as the lead candidate based on its potency and targeting of a conserved sequence proximal to the translational start site. This position was expected to favor the suppression of full-length RAPTOR expression while reducing the potential for residual transcripts to support production of partial RAPTOR protein species. ASO2 treatment produced a dose-dependent reduction in RAPTOR expression and corresponding modulation of mTORC1 pathway activity. Importantly, the ability of these ASOs to engage conserved *Rptor/RPTOR* mRNA sequences across species provides a translational advantage, allowing mechanistic insights from murine disease models to inform therapeutic development for human disease. The identification of additional RAPTOR-targeted ASOs with distinct sequences provides complementary candidates for optimizing the degree and durability of RAPTOR suppression and supports future therapeutic development across mTORC1-driven disorders. More broadly, this work supports the emerging concept that non-enzymatic scaffolding proteins can be effectively targeted using ASO approaches, providing access to signaling nodes that are otherwise difficult to modulate by conventional pharmacological approaches.

Hyperactive mTORC1 signaling is an important pathological mechanism in several genetic epilepsies, subsets of autism spectrum disorder, and multiple cancers (Saxton and Sabatini, 2017). Pathway hyperactivation can arise from mutations in distinct upstream regulators of mTORC1, including components of the GATOR1 and TSC1/2 complexes, as well as from dysregulated metabolic and growth-factor signaling. Extensive work has shown that pharmacological inhibition of mTORC1 with rapamycin or related compounds can be beneficial in TSC models, reducing seizure burden and improving neurological outcomes (Ehninger et al., 2008; Meikle et al., 2008; Sadowski et al., 2022; Zeng et al., 2008). Similar benefits have been reported in GATOR1-deficient models, further supporting mTORC1 as a therapeutic target across these disorders (Ishida et al., 2022; Yuskaitis et al., 2019). Genetic studies further reinforce this model, as partial reduction of RAPTOR through *Raptor* heterozygosity attenuates mTORC1 hyperactivity and improves neurological phenotypes in TSC mutant mice (Karalis et al., 2022). At the same time, the limited efficacy of current anti-epileptic drugs to treat GATOR1-dependent epilepsy, including commonly used sodium channel inhibitors, underscores the need for targeted strategies that address the underlying pathway abnormality across multiple cellular compartments (Wang et al., 2023; Yin et al., 2023). RAPTOR-targeted ASOs offer a precision approach that enables partial reduction of mTORC1 signaling capacity in distinct cellular compartments of the brain while limiting liabilities related to systemic mTORC1 inhibitor drug exposure.

A central finding of this study is the robust *in vivo* efficacy of RAPTOR-targeted ASO treatment across GATOR1-deficient epilepsy models. In both neuronal and astrocytic NPRL2 KO mice, a single ASO2 treatment significantly extended longevity, while longitudinal video monitoring in NPRL2 nKO mice demonstrated sustained suppression of spontaneous seizure activity. Seizure onset in NPRL2 astroKO mice is more variable than in the early-onset NPRL2 nKO model, complicating assessment of treatment effects on seizure activity in small cohorts. However, no seizures were detected after ASO2 treatment in the exploratory NPRL2 astroKO cohort, and future studies incorporating extended video-EEG monitoring may further define the effects of RAPTOR-targeted ASO treatment on seizure activity in this model. The reduced seizure burden, together with the substantial longevity benefit following a single dose, provides evidence that partial RAPTOR reduction with ASOs can produce durable modification of disease-associated phenotypes rather than transient symptomatic control alone. Notably, these therapeutic effects were achieved with only partial RAPTOR reduction, indicating that complete pathway suppression is not required to obtain substantial therapeutic benefit. The durability of ASO2 *in vitro* and *in vivo* is consistent with the prolonged stability of 2’-O-methoxyethyl-modified ASOs in cells and tissues, supporting sustained pharmacological activity within the central nervous system. Importantly, ASO2-mediated reduction of RAPTOR expression did not impair novel object recognition in either control or NPRL2 astroKO mice, supporting preservation of recognition memory following RAPTOR reduction in the central nervous system. Furthermore, brain-directed ASO treatment did not induce hepatic gluconeogenic gene expression characteristic of chronic systemic mTORC1 inhibition under the conditions examined. Consistent with this tolerability profile, rescued NPRL2 nKO mice showed no evidence of leukopenia, granulocytopenia, or anemia, although modest changes in platelet and red-cell indices warrant further evaluation. These findings support the potential of RAPTOR-targeted ASOs for durable modulation of pathological mTORC1 signaling (Backstrom et al., 2024).

Our *in vitro* studies provide additional mechanistic support for the *in vivo* therapeutic efficacy of RAPTOR-targeted ASO2. In primary knockout neuronal cultures derived from NPRL2^fl/fl^ mice, ASO2 normalized pathological network hyperexcitability, as measured by multielectrode array recordings. These findings provide a functional correlate of the seizure suppression observed *in vivo* and support the conclusion that partial RAPTOR reduction is sufficient to normalize neuronal network hyperactivity without broadly suppressing neuronal activity. RAPTOR reduction also corrected astrocytic abnormalities associated with GATOR1 deficiency, including reactive morphology, GFAP expression, Complex II- and IV-linked respiration, mitochondrial ROS, and markers of oxidative damage. Notably, mitochondrial dysfunction has also been reported in TSC patient-derived astrocytes, including reduced respiratory capacity and mitochondrial depolarization, while oxidative stress has been observed in TSC tissue and TSC1-deficient models (Romagnolo et al., 2024; Zimmer et al., 2020). Recent work has further demonstrated that TSC2 loss can drive a cell-autonomous, mTORC1-dependent reactive astrocyte state in human brain organoid models and patient tissue, which can be partially reversed by pharmacological mTOR inhibition (Li et al., 2026). Together with the mitochondrial and redox abnormalities we identified in GATOR1-deficient astrocytes (Hadj-Aissa et al., 2026), these observations suggest that astrocytic dysfunction may represent a convergent feature of pathological mTORC1 activation across distinct mTORopathies. Accordingly, RAPTOR-targeted ASOs may provide a strategy to correct astrocytic dysfunction in TSC and other mTORC1-driven disorders.

The activity of ASO2 in human astrocytes provides an important cross-species extension of these findings. RAPTOR-targeted ASO2 reduced human astrocyte invasion in Boyden chamber assays, demonstrating that it can attenuate an mTORC1-associated cellular phenotype in human cells. This effect is consistent with the broader reduction in astrocytic reactivity observed in ASO2-treated NPRL2 astroKO mice, including reduced GFAP staining and reactive morphology, and supports further investigation of the relationship between RAPTOR-dependent signaling and astrocytic state. Collectively, the correction of disease-associated phenotypes in murine models and modulation of a related cellular phenotype in human astrocytes support the translational potential of RAPTOR-targeted ASOs for neurological disorders characterized by hyperactive mTORC1 signaling.

Our biochemical studies further define how RAPTOR-targeted ASOs can partially restore nutrient-responsive control of mTORC1 signaling despite defective upstream regulation. In NPRL2 KO MEF cells, mTORC1 activity remained elevated during nutrient deprivation, as indicated by persistently elevated phosphorylation of S6K and S6 compared with control cells under EBSS conditions. ASO2-mediated RAPTOR reduction partially restored nutrient sensitivity, enabling suppression of P-S6K and P-S6 toward control levels, consistent with the requirement for RAPTOR to mediate RAG-dependent recruitment of mTORC1 to the lysosomal surface, where its activation occurs (Sancak et al., 2008). Our results therefore suggest that reducing RAPTOR expression limits mTORC1 recruitment and re-establishes nutrient-responsive pathway control despite the absence of GATOR1 function. ASO2 treatment also produced a modest increase in phosphorylated AKT, consistent with relief of mTORC1/S6K-mediated negative feedback on upstream signaling and preservation of mTORC2 activity (Harrington et al., 2004; Sarbassov et al., 2005). Together, these findings support a model in which partial RAPTOR reduction reduces pathological mTORC1 signaling capacity while preserving the ability of the pathway to respond to physiological regulatory inputs.

Collectively, this work establishes RAPTOR-targeted ASOs as a mechanistically distinct strategy for selectively dampening pathological mTORC1 signaling in GATOR1-deficient disorders. By reducing the abundance of RAPTOR, a complex-defining mTORC1 component, this approach lowers pathological signaling capacity without directly inhibiting the mTOR kinase shared by mTORC1 and mTORC2. This provides a distinct route to mTORC1-selective modulation that may avoid liabilities associated with catalytic mTOR inhibition (Feldman et al., 2009; Sevigny et al., 2020; Thoreen et al., 2009). Together, these findings position RAPTOR-targeted ASOs as a precision therapeutic strategy for restoring regulatory control of mTORC1 in GATOR1-dependent epilepsy and related mTORC1-driven disorders.

## Declaration of Interests

P.A.D. and A.D. are inventors on a patent application WO 2026/165665 related to RAPTOR-targeting antisense oligonucleotides described in this study. The intellectual property is owned and managed by Université Laval. All other authors declare no competing interests.

### Declaration of generative AI and AI-assisted technologies in the manuscript preparation process

During the preparation of this work, the authors used Microsoft Copilot for generating the cartoon images for Figure 3A. The authors reviewed and edited the output as needed and take full responsibility for the content of the published article.

## Acknowledgements

This work was supported by grants from the Brain Canada Foundation (Future Leaders in Canadian Brain Research program to PAD), the Canadian Institutes of Health Research (202409PJT-526975 to PAD), Fonds de recherche du Québec – Santé (FRQS J2; Chercheur-Boursier award to PAD), and the Rare Diseases: Models & Mechanisms Network (Fondation du Grand Défi Pierre Lavoie, Genome Canada, and CIHR to PAD). We are grateful to Dr. Benjamin Tu at the University of Texas Southwestern Medical Center for kindly providing the *Nprl2*-floxed mice. We also acknowledge the CERVO Brain Research Centre research platforms and the lab of Martin Lévesque for technical support with neuronal cultures and animal care, and the clinical hematology laboratory at the CHU de Québec-University Laval for blood analysis services. Finally, we thank members of the laboratory for their scientific input, technical assistance, and support throughout this work.

## Methods

### Animal Studies

Generation of the astrocytic and neuronal NPRL2 knockout animals was reported previously (Hadj-Aissa et al., 2026; Hui et al., 2022). Briefly, *Nprl2*^*fl/+*^ mice were crossed with either *VgluT2-IRES*-Cre or *GFAP-IRES*-Cre female mice to generate offspring with heterozygous NPRL2 deletion in glutamatergic neurons or astrocytes, respectively. Subsequent backcrosses with *Nprl2*^*fl/+*^ and *Nprl2*^*fl/fl*^ males were performed, which resulted in control (NPRL2^fl/+^ and NPRL2^fl/fl^) and knockout animals used in this study. Genotypes of all animals were determined by PCR using the following primers: Nprl2 (forward) 5⍰-CTCAGGTTCTACGCAGTGACTTC-31, Nprl2 (reverse) 5⍰-CATGGCGCTGTCTGGATCC-3⍰, Nprl2 (knock-out) 5⍰-CAGGCTTCATACTTCTACCCTC-3⍰, VgluT2 (forward) 5⍰-AAGAAGGTGCGCAAGACG-3⍰, VgluT2 (Cre-reverse) 5⍰-ACACCGGCCTTATTCCAAG-3⍰, VgluT2 (wt-reverse) 5⍰-CTGCCACAGATTGCACTTGA-3⍰, Gfap (forward) 5⍰-TCCATAAAGGCCCTGACATC-3⍰, and Gfap (reverse) 5⍰-TGCGAACCTCATCACTCGT-31. Video recordings were captured using a day/night vision Sony FDR-AX53 camcorder for 24 hours per week while maintaining the mice in a 12h light/dark cycle. Video recordings were manually reviewed for spontaneous seizure-like events, defined by episodes of uncontrolled repetitive twitching or sustained tonic contractions, and the number of affected animals and total seizure episodes were recorded. All animal experiments were approved by the Université Laval Committee on Ethics and Animal Research.

### Antisense Oligonucleotide Screening

A panel of antisense oligonucleotides (ASOs) was designed against conserved mouse and human *Rptor/RPTOR* mRNA sequences, excluding candidates with predicted off-target complementarity using NCBI BLAST analysis. ASOs were synthesized using a gapmer design with 2’-O-methoxyethyl modifications at both termini (Integrated DNA Technologies). The lead candidate, ASO2, had the sequence 5’-CGAGAACCTCCAGCCTT-3’. ASOs were resuspended in ultrapure water for *in vitro* studies or phosphate buffered saline for *in vivo* administration. Initial screening was performed by treating cells with each ASO and assessing RAPTOR protein levels by western blotting at the dose and treatment intervals described in the corresponding figures. A secondary screen was also performed using tiled panels of ASOs surrounding the conserved target sequence, excluding candidates with predicted complementarity to off-target transcripts.

### ASO Administration *in vivo*

For *in vivo* studies, ASOs were prepared in PBS and administered by intracerebroventricular (ICV) or stereotactic intrahippocampal injections, as indicated below. Control animals received either sham injection or an equivalent dose of scrASO2, as specified in the figure legends. Target engagement studies were performed using adult wild-type mice under anesthesia, stereotactically injected at ML ± 1.0mm, AP -0.5mm and DV -2.3mm in each hemisphere with 100 µg/side of ASO. Following recovery, mice were returned to their home cages and maintained. For neonatal survival and efficacy, studies were performed by treating postnatal day 5 (P5) pups with 50 µg /side of ASO using free-hand ICV techniques, adapted from published techniques (Kim et al., 2014). Following injection, pups were returned to the dam and subsequently weaned according to standard breeding procedures. A limited number of peri-procedural deaths occurred within 48 hours of free-hand ICV administration across all cohorts, independent of oligonucleotide sequence, and were excluded from survival analysis. Wet food was supplemented in weaning cages for approximately 2 weeks to ensure nutritional availability. For focal hippocampal studies, 4-month or 5-month old NPRL2 astroKO and littermate control mice were stereotactically injected at LM ± 1.5mm, AP -2.5mm and DV -1.7mm in each hemisphere with 100 µg/side of ASO. Mice were returned to their home cage after surgery and maintained until tissue harvest or downstream analysis, as indicated in the figure legends.

### Novel Object Recognition Test

This test is used to determine recognition memory, as previously described (Pelaez et al., 2023). Briefly, during the first day of testing, 5-month-old mice were placed in the cage and allowed to explore for 5 min to adapt to the new environment. On the second day, two identical objects (familiar objects) were presented to the mice for 5 min. On the third day, corresponding to the test day, the mice were exposed to one familiar and one novel object for 5 min. Interaction time with each object during the duration of the test was recorded and graphed as percentage of total interaction time, defined by nosing, touching, and sniffing.

### Primary Neuron Cultures

Primary cortical neuron cultures were prepared from P0-1 *NPRL2*^*fl/fl*^ mice following previously described methods (Hui et al., 2022). Briefly, dissociated neurons were grown in complete Neurobasal media supplemented with serum-free B-27™ (50:1; Gibco, 17504001), penicillin/streptomycin (50 U/mL; 50 μg/mL; Gibco, 15,140,148) and 0.5 mM L-GlutaMAX (Invitrogen, 35050061). On day-in-vitro 1 (DIV1) cultures were transduced with either control or CRE-expressing lentivirus. At DIV4, cultures were treated with 5 µM Ara-C (Sigma, C1768) and either scrASO2 (control) or RAPTOR-targeted ASO2, as detailed in the figure legend.

### Multielectrode Array (MEA) Recordings

Dissociated primary neuron cultures were generated and plated onto CytoView MEA 24-well plates (Axion Biosystems, M384-tMEA-24W) containing 20,000 cells per well in complete Neurobasal media supplemented with serum-free B-27™ (50:1; Gibco, 17504001), penicillin/streptomycin (50 U/mL; Gibco, 15140148) and 0.5 mM L-GlutaMAX (Invitrogen, 35050061). Spontaneous network activity of the cultures was recorded using a Maestro MEA System (Axion Biosystems) for 5 minutes from DIV6 to DIV21 at 37°C and 5% CO_2_. Data were sampled at 12.5 kHz, digitized, and analyzed using Axion Integrated Studio software with a 200 Hz high-pass and 3 kHz low-pass filter. Burst parameters were set for a minimum of 5 spikes with maximum 100 ms interspike interval. The burst detector was set to detect network burst, with maximum 100 ms interspike interval, minimum 50 spikes, and minimum 35% participating electrodes. After recording, the data were analysed using the Neural Metric Tool and AxIS Metric Plotting Tool modules and graphed using MS Excel.

### Mitochondrial Respiration Analysis

The Oroboros O2K high-resolution respirometer was calibrated in DMEM supplemented with 10% FBS according to the manufacturer’s protocol. Mitochondrial respiration assays were performed as previously detailed (Hadj-Aissa et al., 2026), with the addition of ASO pretreatment. Briefly, control and NPRL3 KO C8-D1A astrocytes were treated with 1 µM of either ASO2 or scrASO2 for 72 hours before analysis, and cell numbers were normalized prior to respiratory measurement. Cells were transferred to the O2k chambers and permeabilized with 100 µg/mL digitonin. Leak respiration was assessed following addition of 5 mM pyruvate, 2 mM malate, and 10 mM glutamate. ADP (2.5 mM) was added to stimulate Complex I-linked respiration, followed by 10 mM succinate to measure combined Complex I- and Complex II-linked respiration. Rotenone (0.5 µM) was added to inhibit Complex I and determine Complex II-linked respiration. Residual oxygen consumption was measured following addition of 2.5 µM Antimycin A and was subtracted from respiratory measurements. Complex IV-linked respiration was assessed following addition of ascorbate and 500 µM N,N,N⍰,N⍰-tetramethyl-1,4-benzenediamine dihydrochloride. Measurements were corrected by subtracting the oxygen consumption rate (OCR) measurement after the addition of 100 mM sodium azide.

### Cell Invasion Assay

NHA cells were trypsinized and seeded onto fibronectin-coated Boyden chamber inserts. Cells were allowed to invade through the matrix-coated porous membrane (5 microns) for 6 hrs, after which they were fixed with paraformaldehyde. Non-invading cells remaining on the upper surface of the membrane were removed before fixation, and cells that invaded through the membrane were imaged at 20X magnification. For each insert, 8x8 tiled fields were acquired and stitched using the Large Image function of NIS-Elements software to generate composite images of 64 fields. Invading cells were quantified from stitched images. Each experimental condition was performed in triplicate. To account for variability in baseline invasion between different experiments, invasion was expressed for each replicate as the ratio of ASO2 to scrASO2-treated controls, and these values were combined across replicate experiments. Because membranes were fixed for imaging, samples from the invasion assay were not suitable for downstream protein extraction.

### Western Blot Analysis

Tissues or cells were homogenized in lysis buffer containing 150 mM sodium chloride, 50 mM sodium fluoride, 100 μM sodium orthovanadate (pH 10.0), 50 mM sodium pyrophosphate tetrabasic, 10 mM β-glycerophosphate, 5 mM EDTA, 5 mM EGTA, and 10 mM HEPES (pH 7.4) and 0.5% Triton X-100 supplemented with complete anti-protease cocktail (Roche). Lysates were pelleted by centrifugation and cleared supernatants, representing the soluble fraction, were boiled in 1x Laemmli sample buffer prior to separation on SDS-PAGE for western blotting using antibodies listed in Table S2.

### Quantitative RT-PCR

Total RNA from tissues or cells was extracted in TRIzol (Invitrogen), following the manufacturers protocol. Primers were selected to span exon-exon junctions when possible. RNA extracts were treated with DNase (Roche) and cDNA synthesis was performed using the High Cap cDNA Reverse Transcription kit (Applied Biosystems). All reactions were performed in triplicate on the QuantStudio 5 Real-Time PCR system (Applied Biosystems). Relative mRNA levels were calculated using the comparative threshold cycle method using U36B4 as the internal control for human or mouse studies, respectively, with QRT-PCR primer sequences listed in Table S1.

### Immunohistochemistry Analysis

Animals were transcardially perfused and cleared with PBS before perfusion with 4% paraformaldehyde (PFA). Brain tissues were collected, fixed overnight in 4% PFA at 4°C, sectioned and processed for immunohistochemistry, as previously described (Hui et al., 2022). Antibodies used for staining tissues are listed in Table S2. Fluorescent images were acquired using identical microscope acquisition settings for all samples to allow for quantitative comparison of signal intensity. *.CZI images were exported from Zeiss Zen as maximum intensity projections in TIFF format to preserve original bit depth and channel information. Quantification was performed using FIJI. To restrict analysis to tissue regions and exclude empty background, a tissue mask was generated from the combined RAPTOR and DAPI signal. Pixel intensity thresholds were applied using an adaptive global threshold using Otsu’s method to identify tissue-containing pixels. Pixels below this threshold were excluded from analysis to remove no-tissue regions and background fluorescence. Regions containing sectioning artifacts (ex. tissue folds) were manually excluded by region prior to quantification to prevent artificial intensity inflation. RAPTOR expression was quantified from the green channel within the tissue mask with measurements normalized to the masked tissue area. Mean and SEM values were calculated across biological replicates within each treatment group.

### MitoSOX Assays

MitoSOX Red assays were performed on cell lines by treating them with 1 µM MitoSOX Red for 15 minutes prior to image acquisition using an epifluorescence microscope according to the manufacturer’s recommendations (Thermo Fisher).

### Statistical Analysis

Statistical analyses were performed using Microsoft Excel and GraphPad Prism 10.4.1. The statistical unit was defined as an independent biological replicate. For comparisons between two groups, Welch’s *t-* tests were used as described in the figure legends. For comparisons involving more than two groups, one-way ANOVA with Tukey’s post-hoc test was used. Survival distributions were compared using the log-rank (Mantel-Cox) test. Data are presented as the mean ± SEM unless otherwise indicated. Statistical significance was defined as *p* < 0.05.

