## Supplemental figures and tables for "RAPTOR-targeted Antisense Oligonucleotides Restrict Pathological mTORC1 Activity in GATOR1-dependent Disease"

**Figure S1. RAPTOR-targeted ASO2 reduces brain RAPTOR expression and dampens mTORC1 signaling *in vivo.* (A) Western blot analysis of RAPTOR expression in brain protein extract from adult mice 14 days after bilateral stereotactic ICV administration of ASO2 (100 µg/hemisphere) or sham treatment. β-actin was used as a loading control. (n=4 mice per group). (B) Quantification of RAPTOR protein abundance from (A), normalized to β-actin and expressed as relative density (R.D.). (C) Representative western blot analysis of RAPTOR, P-S6 (Ser240/244), S6 and β-actin in brain protein extracts from NPRL2 astroKO and control mice treated bilaterally with hippocampal injections of ASO2 (100 µg/hemisphere) or sham at 4 months of age and analyzed one month later. (n=3 mice per group). (D) Quantification of RAPTOR protein abundance from (C), normalized to β-actin and expressed as relative density. (E) Quantification of P-S6 from (C), normalized to total S6 and expressed as relative density. Data are presented as mean ± SEM. Statistical comparisons were performed using two-tailed Welch’s *t*-tests. *a*, p<0.05; *b*, p<0.01; *c*, p<0.001.**

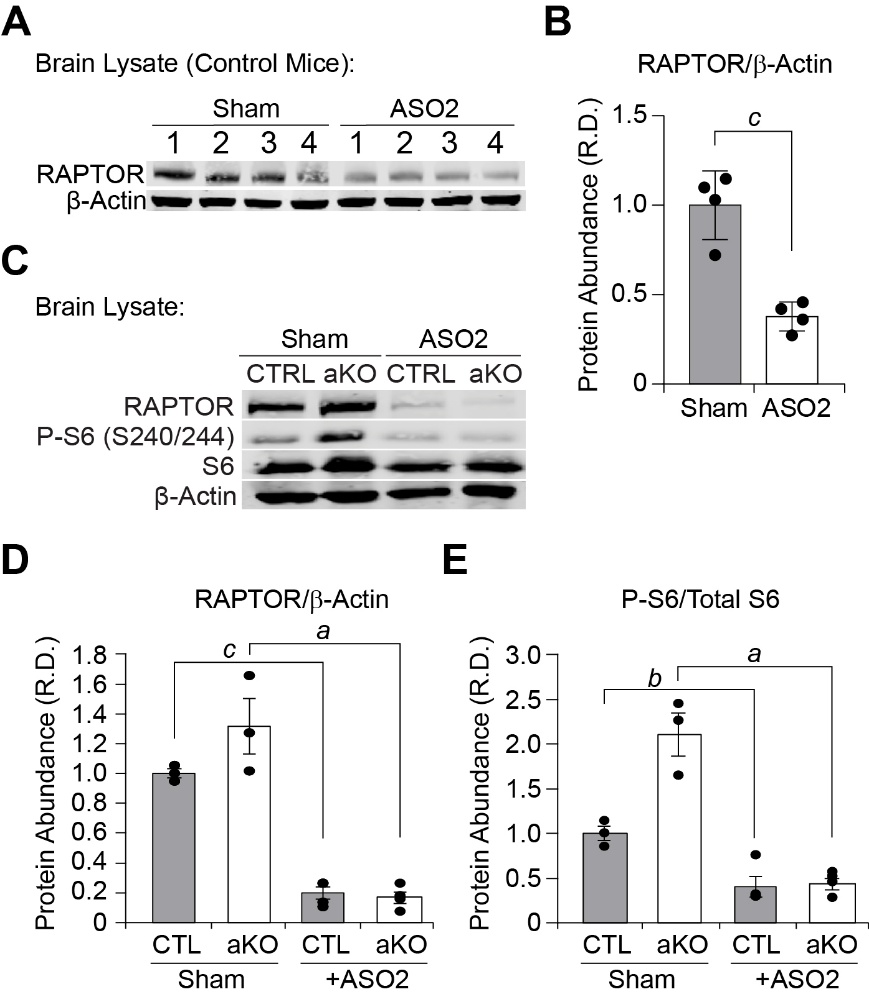

**Figure S2. RAPTOR-targeted ASO2 reduces astrocytic reactivity and oxidative damage in NPRL2 astroKO mice. (A-D) Representative immunohistochemistry and confocal microscopy images of GFAP staining in hippocampal sections from NPRL2 astroKO and control mice treated with ASO2 or sham at 4 months of age and analyzed one month later. GFAP is shown in green and DAPI in blue. Images correspond to the CA1 (A), CA2 (B), CA3 (C) and dentate gyrus (D) regions. (n=3 mice per group). (E) Quantification of GFAP fluorescence intensity within the indicated region from (A-D) expressed in arbitrary units (A.U.). (F) Representative immunohistochemistry and confocal microscopy images of 4-hydroxynonenal (4-HNE) staining in separate sections from the same mice analyzed in (A-D). (G) Quantification of 4-HNE fluorescence intensity from (F). Data are presented as mean ± SEM. Statistical comparisons were performed using one-way ANOVA followed by Tukey’s post-hoc test. n.s., not significant; *a*, p<0.05; *b*, p<0.01; *c*, p<0.001.**

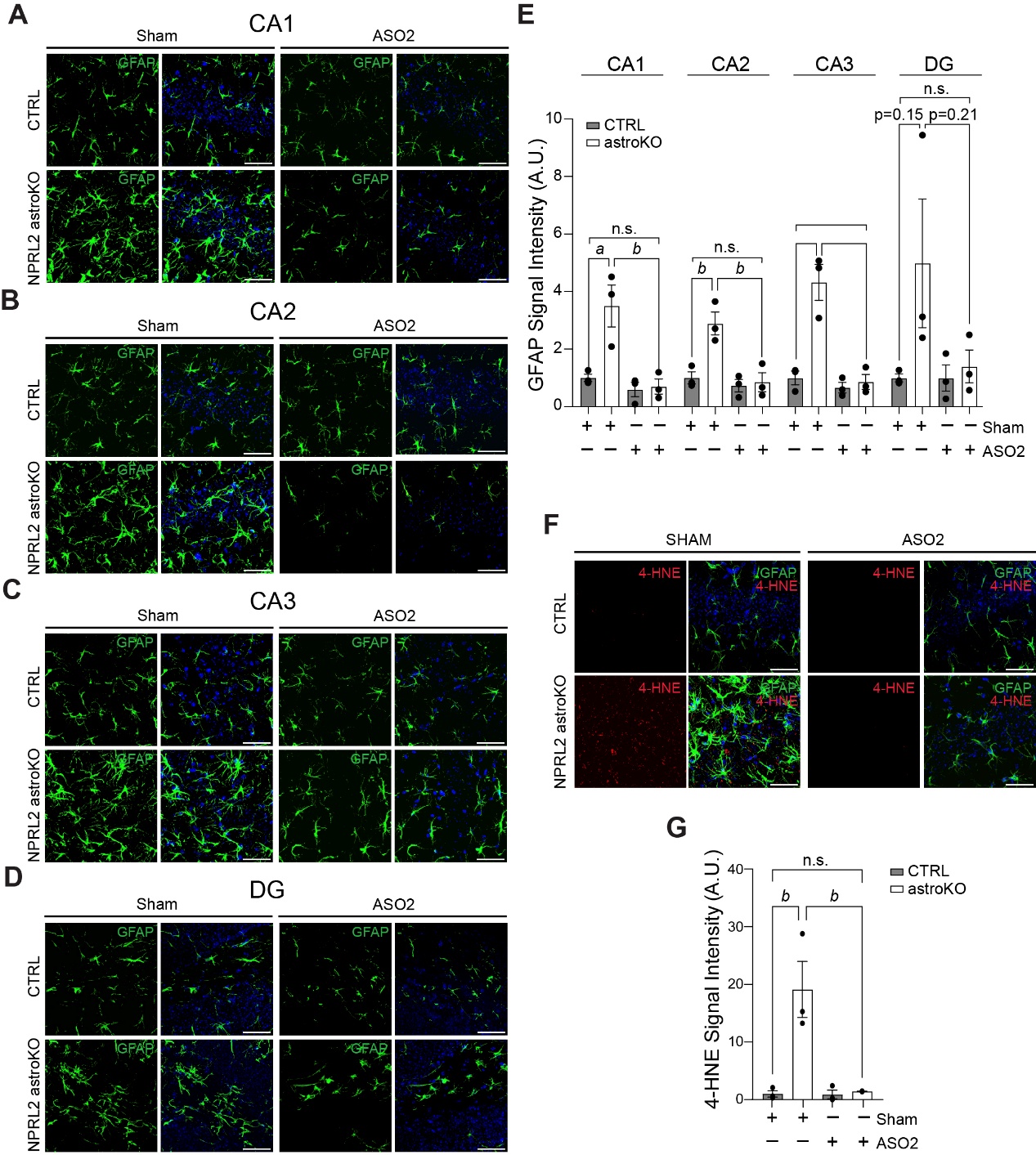

**Figure S3. RAPTOR-targeted ASO2 does not impair recognition memory or increase blood glucose following brain administration.**

**(A) Novel object recognition performance in NPRL2 astroKO and control mice treated with sham, scrASO2 or ASO2 (100 µg/hemisphere) at 4-months old and measured one month after treatment. Statistical comparisons were performed using two-tailed Welch’s *t*-tests. *b*, p<0.01; *c*, p<0.001.**

**(B) Blood glucose measurements in NPRL2 astroKO and control mice following sham, scrASO2, or ASO2 treatment. Glucose levels were measured at one week and one month after treatment using an automated glucometer. No significant increase in blood glucose was observed following ASO2 treatment under the examined conditions. (n=4/6 mice per condition)**

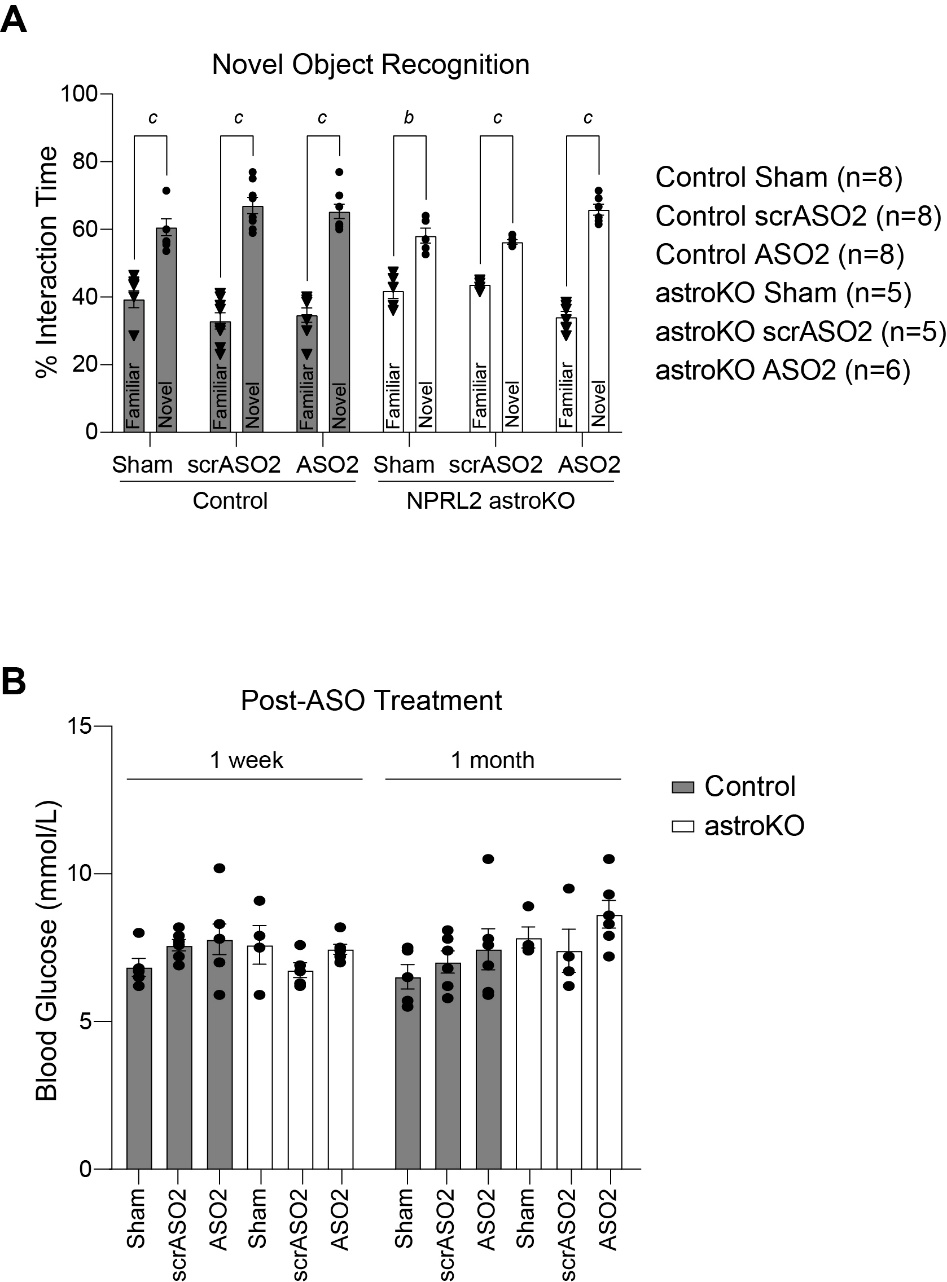

**Table S1.** List of all QRT-PCR primers used in this study.

| **Target** | **Forward** | **Reverse** |
| --- | --- | --- |
| Pepck (mouse) *Phosphoenolpyruvate carboxykinase* | 5’CACCATCACCTCCTGGAAGA3’ | 5’GGGTGCAGAATCTCGAGTTG3’ |
| Rptor (mouse/human) *Raptor* | 5’CTGTCGGCATCTTCCCCTAC 3’ | 5’CACCACCAGCTCGCTGTC3’ |
| U36B4 (mouse) *Ribosomal Protein Lateral Stalk Subunit P0* | 5’CGTCCTCGTTGGAGTGACA3’ | 5’CGGTGCGTCAGGGATTG3’ |
| U36B4 (human) *Ribosomal Protein Lateral Stalk Subunit P0* | 5'CGAGGGCACCTGGAAAAC3' | 5'CACATTCCCCCGGATATGA3' |

**Table S2.** List of all antibodies used in this study.

| **Target** | **Source** | **Catalog Number** |
| --- | --- | --- |
| 4-HNE | Thermo Fisher Scientific | MA5-27570 |
| β-actin | CST | 3700S |
| GFAP | Millipore Sigma | AB5541 |
| P-S6 (S240/244) | CST | 2215S |
| RAPTOR (monoclonal) | CST | 2280S |
| RAPTOR (polyclonal) | Invitrogen | 42-4000 |
| S6 | CST | 2217S |
| P-AKT (Ser473) | CST | 4060S |
| AKT | CST | 2920S |
| P-S6K (T389) | CST | 9205S |
| S6K | CST | 9202S |
| Secondary: Alexa Fluor 488 Goat anti-Chicken | Thermo Fisher Scientific | A32931 |
| Secondary: Alexa Fluor 546 Goat anti-Mouse | Thermo Fisher Scientific | A-11030 |
| Secondary: Alexa Fluor 647 Goat anti-Rabbit | Thermo Fisher Scientific | A-21245 |
| Secondary: Alexa Fluor 647 Goat anti-Guinea Pig | Thermo Fisher Scientific | A-21450 |
| Secondary: IRDye 680RD Goat anti-Mouse IgG | LICORbio | 926-68070 |
| Secondary: IRDye 800CW Goat anti-Rabbit IgG | LICORbio | 926-32211 |
